# Insights on the mutational landscape of the SARS-CoV-2 Omicron variant

**DOI:** 10.1101/2021.12.06.471499

**Authors:** Nathaniel L. Miller, Thomas Clark, Rahul Raman, Ram Sasisekharan

## Abstract

The SARS-COV2 Omicron variant has sparked global concern due to the possibility of enhanced transmissibility and escape from vaccines and therapeutics. In this study, we describe the mutational landscape of the Omicron variant using amino acid interaction (AAI) networks. AAI network analysis is particularly well suited for interrogating the impact of constellations of mutations as occur on Omicron that may function in an epistatic manner. Our analyses suggest that as compared to previous variants of concern, the Omicron variant has increased antibody escape *breadth* due to mutations in class 3 and 4 antibody epitopes as well as increased escape *depth* due to accumulated mutations in class 1 antibody epitopes. We note certain RBD mutations that might further enhance Omicron’s escape, and in particular advise careful surveillance of two subclades bearing R346S/K mutations with relevance for certain therapeutic antibodies. Further, AAI network analysis suggests that the function of certain therapeutic monoclonal antibodies may be disrupted by Omicron mutations as a result of the cumulative indirect perturbations to the epitope surface properties, despite point-mutation analyses suggesting these antibodies are tolerant of the set of Omicron mutations in isolation. Finally, for several Omicron mutations that do not appear to contribute meaningfully to antibody escape, we find evidence for a plausible role in enhanced transmissibility via disruption of RBD-down conformational stability at the RBD-RBD interface.

## Introduction

The SARS-CoV-2 Omicron (B.1.1.529) variant of concern (VOC) has now been detected globally, seizing the world’s attention due to preliminary data suggesting enhanced transmission and reinfection as compared to the Delta variant [Pulliam 2021]. Several of omicron’s spike mutations have been observed in other VOC and demonstrated to enhance transmissibility and confer varying degrees of escape from neutralizing antibodies [Liu 2021a, Liu 2021b, Ku 2021, Zhou 2021]. However, numerous Omicron mutations have not been observed on previous VOC nor characterized rigorously in terms of their functional effects. Still, the position of many unknown Omicron mutations within antibody epitopes has prompted concerns that the efficacy of vaccines and therapeutic antibodies could be significantly reduced against Omicron, prompting to policy decisions and research prioritizations with far reaching consequences.

In this study, we analyze the receptor binding domain (RBD) mutational landscape of the Omicron VOC using amino acid interaction (AAI) networks. AAI network analysis is particularly well-suited for understanding the impact of constellations of mutations residing within and adjacent to an antibody epitope as occurs on the Omicron variant. For example, AAI network analysis considers how mutation of a residue that does not directly interact with a given antibody paratope (e.g., via a hydrogen bond between side chain and antibody CDR) may still perturb antibody binding if the residue plays a significant structural role in supporting other sites that interact directly with the antibody. AAI networks quantitate these indirect relationships through which Omicron’s mutation constellation may substantially perturb the chemical and physical properties of RBD epitope surfaces. We apply the AAI network lens to describe the potential impact of Omicron RBD mutations on polyclonal antibody responses and therapeutic monoclonal antibodies. We further discuss the limitations in interpreting efficacy of therapeutic antibodies based on existing *in vitro* data of isolated mutations. Finally, we present possible functional roles for Omicron mutations that are not predicted to substantially enhance antibody evasion. Our analysis suggests these mutations may modulate the energetics of the RBD-up transition toward enhanced infectivity.

## Results

### The impact of Omicron mutations on polyclonal antibody evasion

Toward investigating the antigenic impact of the Omicron RBD mutations, we first mapped RBD antibody epitopes using an AAI network technique [Soundararajan 2011, Miller 2021]. Our RBD epitope map includes at least 10 antibodies from each of the four structural classes of anti-RBD antibodies [Barnes 2020] and thus represents the major functional components of population-level polyclonal antibody responses [Hastie 2021].

Using the RBD epitope map we first examine Omicron escape *breadth* across the polyclonal sera response, finding that Omicron mutations occur at RBD sites that interact with all antibodies examined and spanning the four antibody classes (**Figure 1**). In contrast, the RBD mutations of the Beta and Delta variants are confined to sites within class 1 and 2 antibody epitopes, with the exception of Beta N501Y, which interacts indirectly with certain class 3 antibodies. This is consistent with experimental evidence documenting Beta escape from class 1 and 2 antibodies [Wibmer 2021, Yuan 2021] and Delta escape primarily from class 2 antibodies [Cheng 2021]. Our analysis therefore suggests that Omicron variant may have increased antibody escape breadth as compared to previous variants, and that this breadth is driven by mutations in class 3 and 4 antibody epitopes.

**Figure 1:**
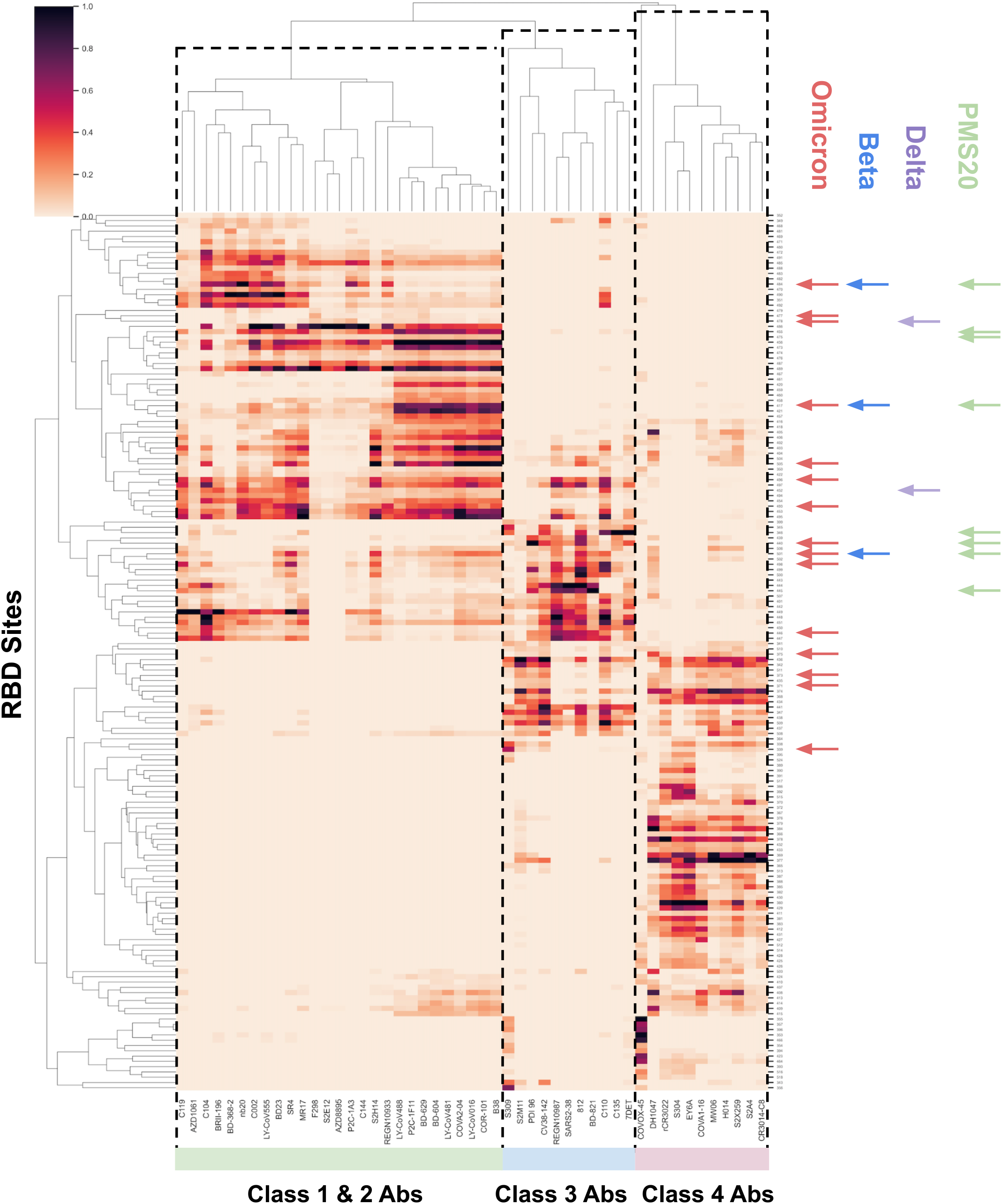
RBD epitopes and variant mutational constellations. AAI networking between a panel of antibodies and nanobodies covering all anti-RBD antibody classes (x-axis) and RBD sites (y-axis) is shown, with networking strength annotated as heat map intensity. RBD sites mutated on the Omicron, Beta, Delta, and PMS20 RBDs are highlighted by red, blue, purple, and green arrows, respectively. The Beta and Delta variant mutations primarily reside at sites corresponding to class 1 and 2 antibodies, the PMS20 mutations occur at sites residing within class 1-3 epitopes, and the Omicron mutations cover the epitopes of all four antibody classes. Further, Omicron and PMS20 feature several class 1 and 3 antibody mutations suggestive of escape depth in these classes. The four Omicron mutations affecting class 4 epitopes do so via indirect networking, though may still cumulatively affect antibody binding at this epitope region.

Class 3 antibodies are potent neutralizers that are immunodominant for certain individuals [Greaney 2021], while class 4 antibodies tend to be weakly neutralizing [Barnes 2020] and thus play a lesser role in escape from polyclonal sera responses. Our network analysis ranks Omicron mutations N440K, G446S, G496S, and Q498R as most likely to confer enhanced class 3 antibody escape based on these sites having the strongest network interactions with the antibodies surveyed. The RBD of PMS20, a variant that was designed to escape neutralization from most convalescent and polyclonal sera, features similar class 3 mutations to Omicron at sites 440 and 445, yet also features an R346K mutation that Omicron lacks [Schmidt 2021]. R346 is the most strongly networked residue for certain class 3 antibodies in our analysis including C135, suggesting PMS20 could escape more effectively from the class 3 antibody component of sera than Omicron.

The Omicron mutations occurring in class 4 antibody epitopes (G339D, S371L, S373P, S375F) are predominantly indirectly networked to class 4 antibodies. While indirect networking would typically suggest that these mutations are unlikely to confer significant class 4 antibody escape in isolation, it is plausible that the combined indirect effects of these four mutations—which confer major chemical changes—could meaningfully alter the local structure in this RBD region and thus perturb class 4 antibodies. Still, Omicron lacks mutation at a site that is strongly directly networked to multiple class 4 antibodies such as 369, 377, 378, or 384.

Further, we note the that Omicron mutations may also enhance escape *depth* from class 1 antibodies beyond that observed for the Beta variant due to accumulation of additional mutations in class 1 antibody epitopes beyond the shared Beta/Omicron mutation K417N. This feature could have contributed to PMS20’s escape from convalescent and vaccinee sera [Schmidt 2021], as it can be seen in **Figure 1** that PMS20 also features multiple class 1 escape mutations. The polyclonal antibody response to infection and vaccination has been shown to broaden over the months following exposure due to persistent somatic mutation of antibody CDRs [Wang 2021]. This broadening process can increase an antibody’s tolerance of escape mutations within its epitope [Muecksch 2021], leading to polyclonal responses that became better at neutralizing the Beta variant over time. We find that Omicron has accumulated multiple tightly clustered mutations and therefore may have enhanced escape from matured polyclonal responses that are tolerant of certain class 1 antibody escape mutations such as K417N and N501Y (see **Figure 1**). Specifically, Omicron mutations at residues Q493, G496, Q498, and Y505 cluster closely with the Beta and Omicron escape mutations at residues K417 and N501 indicating escape depth from broad class 1 antibody responses. Notably, however, PMS20 was not able to fully escape from polyclonal responses generated from infection followed by infection [Schmidt 2021], suggesting that Omicron’s class 1 escape depth alone may not confer this ability.

### Omicron mutations and their effects on therapeutic antibodies

We subsequently applied AAI networks to examine the Omicron mutations in the context of their ability to evade neutralization by therapeutic antibodies of current clinical relevance (**Figure 2**). An important perspective offered by our network approach is the impact of mutations that are not directly located at the interface of antibody-antigen complexes on enhancing or disrupting their interaction. Specifically, it is important to consider the allosteric effects of the many Omicron mutations and how they may cooperatively affect antibody binding to the Omicron RBD by modulating the structural and chemical features of the epitope surface. Our results present an accessible and concise visualization of the network interactions between RBD sites mutated on Omicron and authorized therapeutic antibodies.

**Figure 2:**
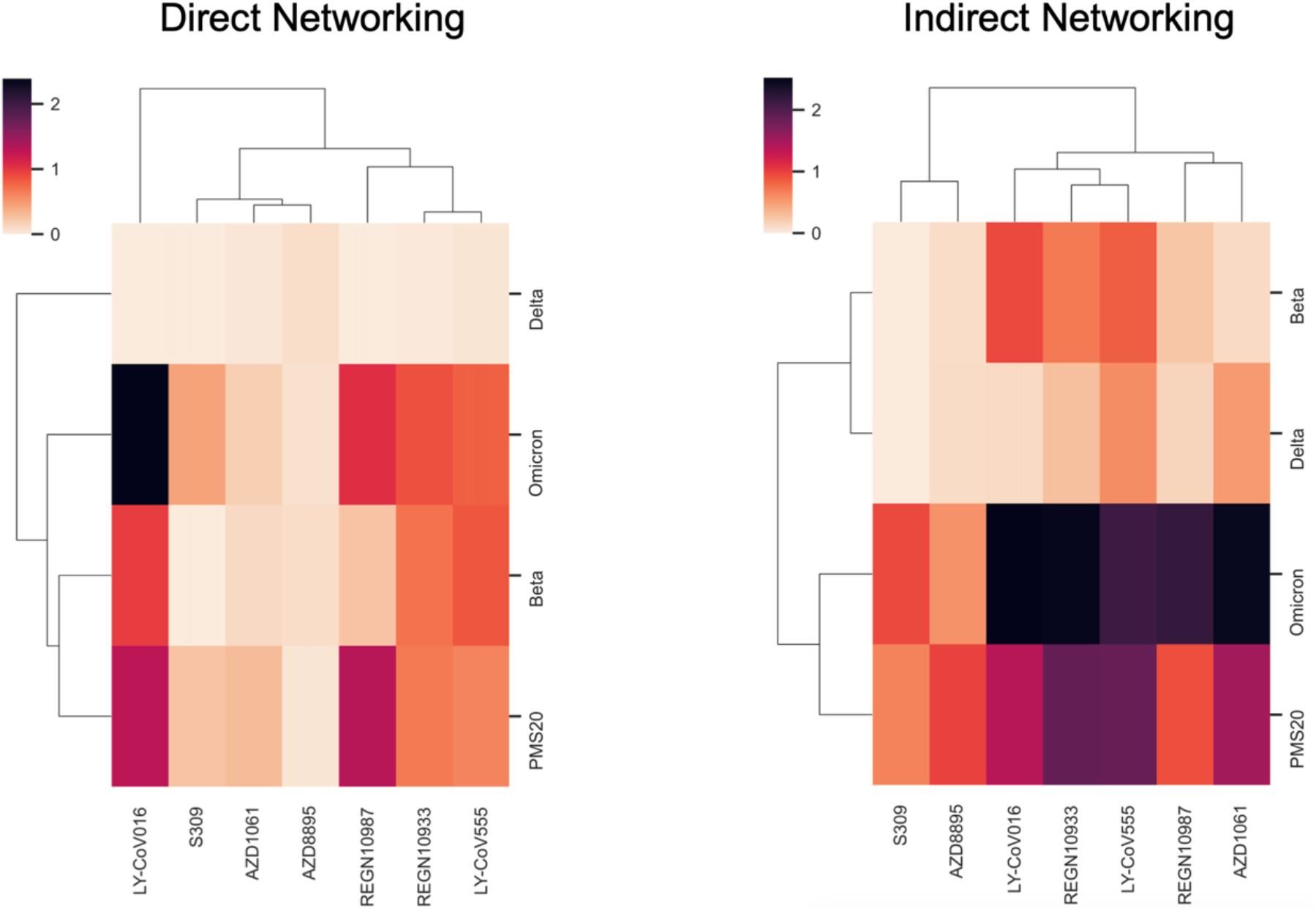
The Omicron, Beta, Delta and PMS20 RBDs versus therapeutic mAbs. The data in Figure 1 is refined to highlight the cumulative direct and indirect networking between the sites of Omicron mutations and the currently authorized therapeutic mAbs. The cumulative networking between these antibodies and the Beta, Delta, and PMS20 RBDs is shown for comparison. AAI network diagrams describing connections between specific sites are provided in the supplement.

We find that Omicron mutations occur at sites that are strongly directly networked to REGN10987+REGN10933 (*Casirivimab* + *imdevimab*; sites 417, 440, 446, 484, 493, 496, 498) and LY-CoV016 + LY-CoV555 (*Bamlanivimab* + *etesevimab*; sites 417, 484, 493, 501, 505), which suggests the binding event of these antibodies will be disturbed by Omicron mutations. In contrast, Omicron mutations do not appear to strongly *directly* interact with *Sotrovimab* (S309) or *Tixagevimab* + *Cilgavimab* (AZD7442 + AZD8895). Our findings on the basis of direct networking between epitope residues in Omicron variant and paratope residues in the neutralizing antibodies align with existing analyses and commentary for these therapeutic antibodies derived from mutagenesis screens based on the binding perturbation induced by the Omicron mutations *individually and in isolation* [Regeneron, Cathcart 2021, Greaney 2021].

However, when further considering the cumulative indirect effects of the Omicron mutations, we observe strong indirect networking from the mutated Omicron sites to AZD1061 and moderate indirect networking to AZD8895 and S309. Examining the S309 AAI network in closer detail (**Figure S1**) shows indirect interactions between S309 and the Omicron mutated sites 339, 373, 440, and 446. Site 339 networking, however, drives the majority of the total indirect networking between the Omicron mutated sites and S309. A recent study suggesting that S309 is tolerant of G339D in isolation is therefore reassuring [Cathcart 2021]. Still, as four Omicron mutations may indirectly alter the properties of the S309 epitope surface, it is important to verify S309’s binding and neutralization in the context of RBD bearing all Omicron mutations. Further, since the submission of this manuscript, Cathcart et al., updated their preprint with pseudovirus neutralization data in the context of the full set of Omicron mutations, recording a <3-fold reduction in neutralization. This reduction is consistent with the AAI model prediction that the cumulative indirect effects of the Omicron mutations moderately perturb the S309 binding surface. Building on this observation, another important residue position in the epitope surface of S309 is R346. The AAI analysis shows that R346 is networked both directly and indirectly to the S309 paratope in the context of the other Omicron mutations. While isolated mutations at R346 only minorly impact S309 binding and neutralization [Starr 2021, Cathcart 2021], the AAI network suggests R346 mutations are likely to have a greater impact in the context of the other Omicron mutations. Several Omicron subclades have been detected with R346S and R346K mutations. Therefore, it is important to monitor if there is a further reduction in the susceptibility (>5-fold change as defined in the *Sotrovimab* EUA Fact Sheet) of subclades with the R346 mutations to neutralization by S309.

Similarly, such indirect effects may impact the recently identified anti-RBD antibodies such as ADG-2 that target the conserved epitope shared across clade 1a and 1b coronaviruses [Rappazzo 2021]. The AAI network model indicates that Omicron mutations at sites 496, 498, 501, and 505 are networked to the critical epitope residues D405, G502, G504, and Y505 (**Figure S2**). Rappazzo et al., further reported that Y505C/N/S knocked down ADG-2 binding potency. Single point mutation analysis on an epitope does not consider the extent to which mutations at proximal residues may impact the epitope-paratope binding interaction. For example, ADG-2 does not bind the RaTG13 RBD, and RatG13 shares sequence identity with Omicron at the four critical ADG-2 binding sites 405, 502, 504, and 505, demonstrating the importance of the network context of these four critical sites.

### Potential additional functional roles of Omicron RBD mutations

Given the substantial apparent transmission advantage of Omicron over Delta, it is plausible that certain Omicron RBD mutations contribute to enhanced transmission by mechanisms other than or in addition to antibody escape. Above (**Figure 1**), we identified several Omicron mutations that did not appear to play a major role as polyclonal epitope constituents or whose predicted contribution to antibody evasion was unlikely to confer a substantial fitness advantage due to the mutations occurring in class 4 antibody epitopes. Here, we highlight two mechanisms by which these RBD mutations may contribute to Omicron’s fitness via enhanced transmissibility.

#### Enhanced ACE-2 binding

There is existing evidence that certain Omicron mutations enhance ACE-2 binding and this is a leading hypothesis to explain Omicron’s enhanced transmission. In particular, Zahradník et al., identified enhanced ACE2 binding by RBD’s bearing Q498R and S477N when combined with the N501Y mutation [Zahradník et al]. Importantly, these mutation combinations resulted in a synergistic effect extending beyond the increased ACE-2 binding predicted when the effect of these mutations assayed individually was summed. It is therefore likely that a similar synergy effect occurs on the Omicron variant at these three sites and may also include additional Omicron mutations. Using AAI networks to examine the local residue dependencies in this vicinity, we observe a network extending from the known synergistic pair 498+501 to position 505, suggesting Omicron H505 may modulate the synergistic ACE-2 binding effect for Q498R + N501Y.

#### RBD-RBD Interface Stability

While Omicron’s class 4 mutations may result in antibody escape via indirect and cooperative mechanisms, evolution of such a triplet mutation toward this escape function is not parsimonious. Indeed, other sites in class 4 epitopes appear to confer antibody escape with just single mutations. We therefore suspect that mutations S371L, S373P, and S375F may provide a fitness advantage other than escape. In a literature review, we found that Wrobel et al., had previously identified a role for sites S371 and S373 in stabilizing the RBD-RBD interface in the spike RBD-down conformation [Wrobel 2020]. Further, Wrobel et al., identified interactions between sites 369, 371, 373, 403, 440, 493, and 505 as possibly explaining differences between SARS-CoV-2 and RaTG13 in RBD-RBD interface stability with implications for the RBD-up transition that enables ACE-2 binding and infection. Remarkably, Omicron mutations occur in five of these seven sites. Additionally, mutations such as D614G have previously been linked to increased infectivity via destabilization of the RBD-down spike protein conformation [Berger 2020].

We therefore built a homology model of the Omicron RBD-RBD interface and investigated the influence of the Omicron mutations on interface stability (**Figure 3**). We find that for the specific interface interaction highlighted by Wrobel et al., consisting of sites 373, 403, and 505, the Omicron mutations appear to disrupt hydrogen bonding that exists for the wild-type RBD from Y505 to S373 and Y505 to R403. The loss of these bonds is reflected in the positive ΔΔG for Y505H and S373P, yet partially compensated for by a more energetically favorable R403 environment. A large and positive ΔΔG is also observed upon mutation at site 375. Further, for our Omicron model the RBD-RBD interface buried surface area decreases by 40 Å^2^ and the interface surface complementarity decreases from 0.60 to 0.54. The potential for a significant perturbation at this interface is also captured by the AAI network analysis, which identifies numerous direct and indirect interactions from sites 371, 373, 375, 440, 493, 505 extending across the RBD-RBD interface. Together, these data suggest the Omicron mutations may destabilize the RBD-RBD interface in the RBD-down spike conformation, which may facilitate transition to the infectious RBD-up state. Interestingly, such a functional change may also bear a tradeoff with increased susceptibility to antibody neutralization via increased surface exposure of the RBD neutralizing epitopes. Indeed, these predictions can be validated experimentally and explained in more mechanistic detail once an Omicron spike structure is solved.

**Figure 3.**
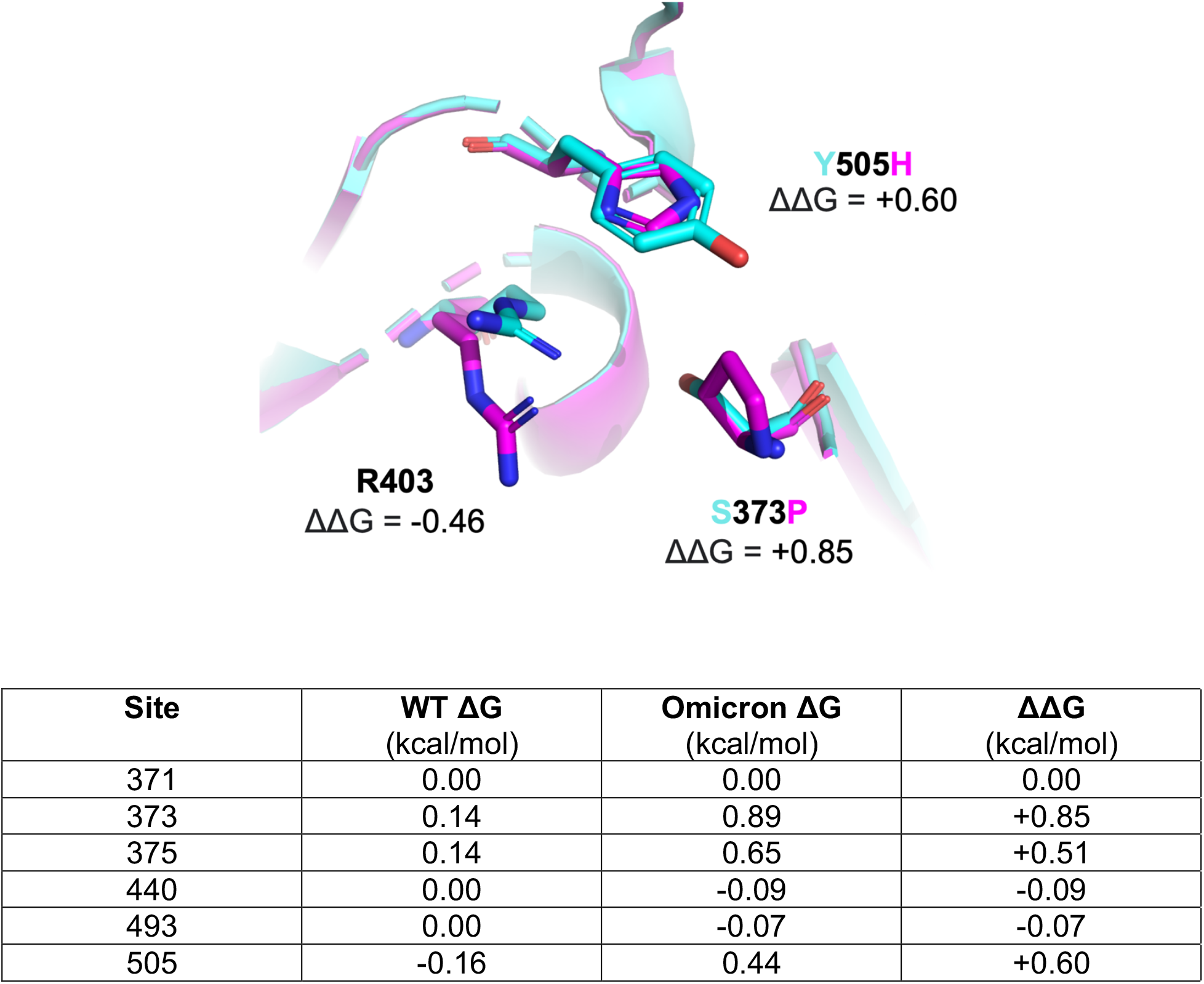
The effects of Omicron mutations on the RBD-RBD interface. **Top**, an Omicron RBD-RBD homology model overlayed with the wild-type RBD-RBD interface in the furin-cleaved state. Omicron mutations S373P and Y505H reduce the energetic favorability of the RBD-RBD complex at these sites, though enable a more energetically favorable conformation for the unmutated R403. **Bottom**, the energetic contribution of all RBD-interface sites mutated on Omicron to RBD-RBD complexation.

## Discussion

The network-based structural analysis of the mutational constellation of the Omicron variant presented here describes Omicron’s escape breadth and highlights key questions for investigation. We find evidence for enhanced escape breadth of Omicron as compared to the Beta and Delta variants due primarily to class 3 escape mutations as well as increased escape depth within class 1 epitopes. We further identify plausible class 4 antibody escape via the mutations at sites 371, 373, and 375 which may alter class 4 epitope surfaces indirectly, but note that escape from class 4 antibodies is unlikely to confer a meaningful escape fraction from sera and additionally that these sites are not the dominant class 4 escape sites.

At this time, it is difficult to predict the extent to which Omicron’s increased escape breadth at class 3 and 4 antibody epitopes and escape depth at class 1 antibody epitopes will read through to increased reinfection of convalescent and vaccinated individuals. Importantly, although neutralizing antibodies are correlated with vaccine protection against infection [Khoury 2021], vaccine efficacy against severe disease is likely to be preserved even in the case of complete antibody escape due to non-neutralizing antibodies and T cell responses [Cevik 2021]. Still, study of the synthetic RBD construct PMS20 [Schmidt 2021] which bears a similar mutational constellation to Omicron offers clues as to potential Omicron antibody evasion. PMS20 escapes neutralization by the majority of convalescent and vaccinated individuals yet cannot fully escape from sera from individuals that are infected and then vaccinated.

Further, PMS20 bears a previously identified escape mutation at site 346 (class 3 and 4 escape) that Omicron does not, suggesting that PMS20 may evade a component of the polyclonal antibody response that Omicron is currently susceptible to. Notably, clade of Omicron have the identified bearing R346K (15 descendants at the time of this writing) and R346S (5 descendants) and should be monitored closely [Hadfield 2019, Shu 2017]. As described earlier, Omicron bearing R346 mutations may also impact the susceptibility of the virus to neutralization by therapeutic antibodies. In particular, R346 mutations may reduce the susceptibility of the virus to Sotrovimab.

Further, it will be important to determine if the accumulation of class 1 escape mutations, which provide escape depth, reads through to a meaningful escape impact or reduction in vaccine efficacy. Thus far, such depth seems to provide little additional escape benefit given that the combination of K417 and N501 mutations appears sufficient to escape from class 1 antibodies in most convalescent and vaccine sera [Yaun 2021]. However, it is notable that the PMS20 variant was unable to fully escape from polyclonal responses generated by infection followed by vaccination [Schmidt 2021]. Such “hybrid” immunity provided exceptionally potent and broad polyclonal responses [Crotty 2021]. It is plausible Omicron’s additional class 1 antibody escape mutations provide a benefit against such responses, which could suggest these Omicron mutations represent multiple rounds of virus-host evolution. Teasing out these specifics of Omicron’s escape breadth and depth will be particularly informative for designing vaccine boosters resilient to constellations of escape mutations both across antibody classes as well as within immunodominant class 1 & 2 epitopes.

Previous variants presented with mutations on largely orthogonal epitope surfaces targeted by distinct antibody classes, yet Omicron’s RBD mutational landscape is highly clustered within overlapping antibody epitopes. Throughout this study we therefore highlight the importance of considering the complex structural and chemical relationships between combinations of mutations within or adjacent to antigenic surfaces. While single mutation escape studies are unlikely to provide a complete mapping of how mutation constellations may perturb antibody binding, our AAI network approach quantitates these complex relationships between proximal residues and interface structure. The direct component of our network analysis was highly consistent with predictions on therapeutic antibody escape provided by traditional single mutation predictions. In contrast, the indirect component identified a number of allosteric network interactions between Omicron mutation sites and key antibody binding residues for S309 and ADG-2. Experiments in which the Omicron mutation constellation is present in full can validate whether such indirect effects sufficiently modulate the properties of the epitope surface to impair binding.

Finally, we find evidence in our structural analysis and in the literature for mechanisms other than immune escape by which Omicron mutations enhance fitness. While mutation combinations such as Q498S and N501Y have been previously shown to epistatically enhance ACE-2 binding, our network analysis suggests that Omicron H505 may also alter the synergistic behavior between these two sites. Further, based on the previous work of Wrobel et al. and our Omicron RBD homology model, we present evidence of a potential role for Omicron mutations in destabilizing the RBD-RBD interface to increase the propensity for the RBD-up conformation. Increased occupancy of the RBD-up state may also present a fitness tradeoff resulting from greater surface exposure of the RBD antibody epitopes. Yet, Omicron’s antibody evasion may partially negate this penalty.

In the probable event of continued Omicron ascent across the globe, determining the various functional roles of the Omicron mutations will be critical. In particular, the complex relationships between Omicron mutations and their contribution to antibody escape and enhanced transmissibility. Network-based approaches such as those presented in this study are particularly well suited for modeling such indirect interactions.

## Acknowledgements

NLM was supported in part by T32 <u>ES007020</u>/ES/NIEHS NIH and by SMART, Singapore.

## Author Contributions

N.L.M. and T.C. conducted the analyses; all authors designed the analyses and wrote the paper.

## Methods

### Amino Acid Interaction (AAI) Network

For each antibody-antigen complex, AAI network scores between every pair of residues within the structure were computed as described previously [Soundararajan 2011, Miller 2021]. Briefly, network scores consider all contacts between both side-chain and backbone atoms, and weight these contacts according to the energetics of the expected interaction. Interactions examined include hydrogen bonds, pi bonds, disulfide bonds, polar interactions, salt bridges, and Van der Waals interactions. Given that the SIN computation is weighted toward side chain interactions, glycine networking was interpolated from nearby residues to estimate the local effect of glycine mutations. The following metrics were defined and computed for all surveyed structures from the resulting interaction network. Direct Networking: For a given residue on RBD, the sum of all interactions between the RBD residue and all residues on the complexed antibody/nanobody. Indirect Networking: For a given residue on RBD, the sum of all interactions between the RBD residue and all other residues on RBD which are directly networked to an antibody/nanobody paratope. Total Networking: For a given residue on RBD, the sum of direct and indirect paratope networking scores. Note that for all networking metrics, scores are normalized to the highest networking score within each RBD-mAb complex. The following structures were analyzed: 6XC7, 6XDG, 6ZER, 7B3O, 7BEL, 7BWJ, 7BYR, 7BZ5, 7C01, 7C8V, 7C8W, 7CAH, 7CDI, 7CDJ, 7CH4, 7CH5, 7CHH, 7DET, 7DPM, 7EY5, 7EZV, 7JMO, 7JMW, 7JVA, 7JVB, 7JW0, 7JX3, 7K43, 7K45, 7K8S, 7K8U, 7K8V, 7K8W, 7K8Z, 7K90, 7K9Z, 7KMG, 7KMG, 7KMH, 7KMI, 7KZB, 7L7E, 7LD1, 7LM8, 7MKM, 7MZK, 7RAL.

### AAI Interaction Matrix, Clustering, and Visualization

mAb-epitope clustering: Direct, Indirect, and Total networking scores for each residue across all mAb and nanobody complexes were plotted using clustermap from the Seaborn statistical data visualization package [Waskom 2021]. AAI networks were visualized using NetworkX [Hagberg 2008].

### Omicron RBD-RBD Interface Analysis

A homology model of the Omicron RBD mutations was built using SWISS-Model [Waterhouse 2018] and the 6ZGI [Wrobel 2020] template. Subsequently, all side chains were repacked in PyRosetta [Chaudhury 2010] for the homology model as well as for wild-type control. Finally, the Omicron and WT RBD-RBD interfaces were analyzed and compared using PDBePISA [Kirssinel 2007]. Surface complementarity was computed using the InterfaceAnalzyerMoveer in PyRosetta [Chaudhury 2010]. Interfaces visualization were generated using PyMOL [PyMOL].

## Supplementary Figures

**Figure S1:**
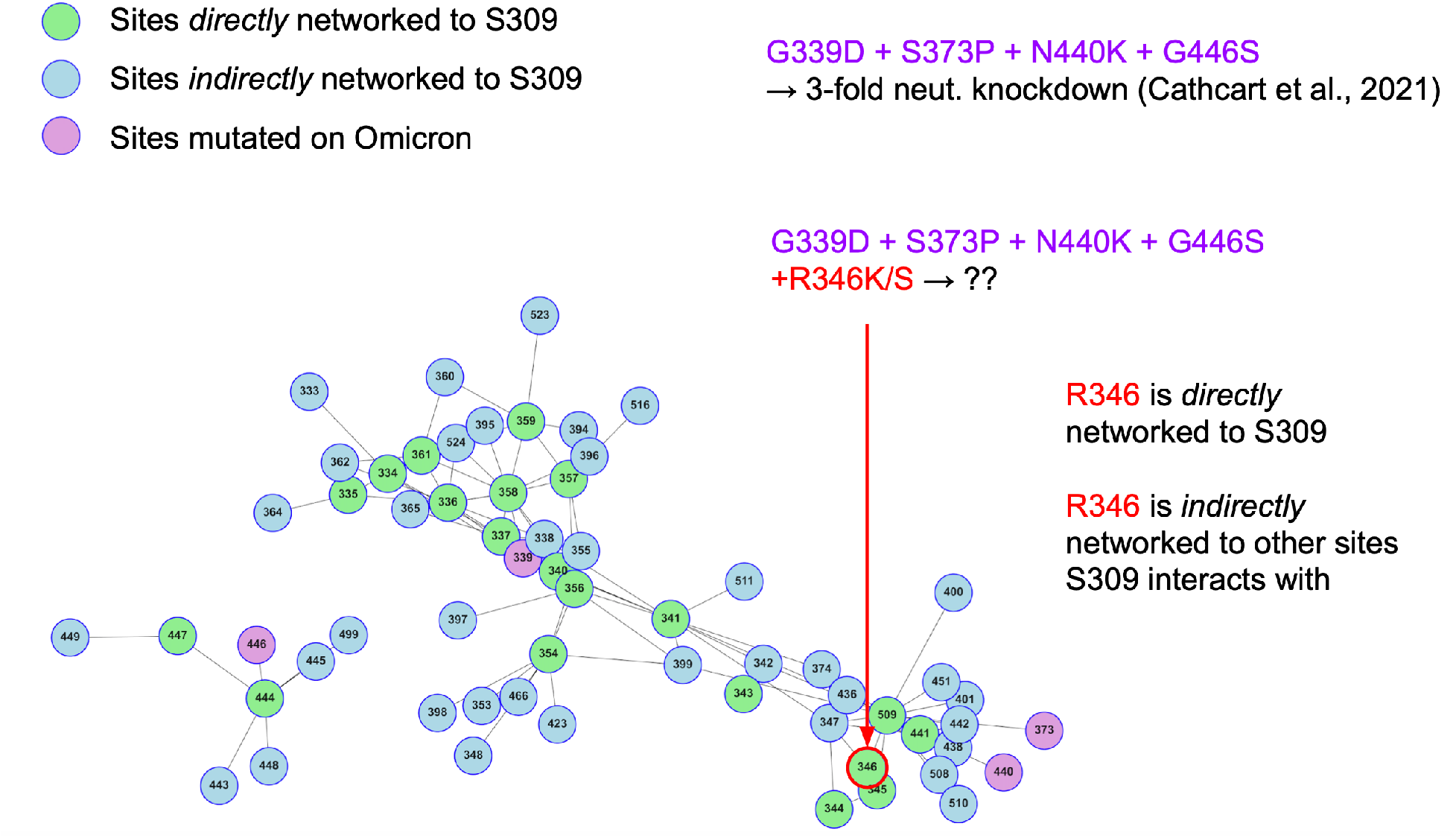
Indirect AAI network of the S309 epitope. RBD sites that directly interact with S309 are shown in green. RBD sites that indirectly interact with S309 are shown in blue and purple, where purple annotates sites mutated on the Omicron variant. Residue R346, which is mutated to S or K in certain Omicron sub-clades, is highlighted in red. Edges between notes are weighted according to network interaction strengths, such that the Euclidean distance between nodes maps inversely to the networking strength between the two sites on RBD. Adjacent node pairs are therefore more strongly networked together than distant node pairs.

**Figure S2:**
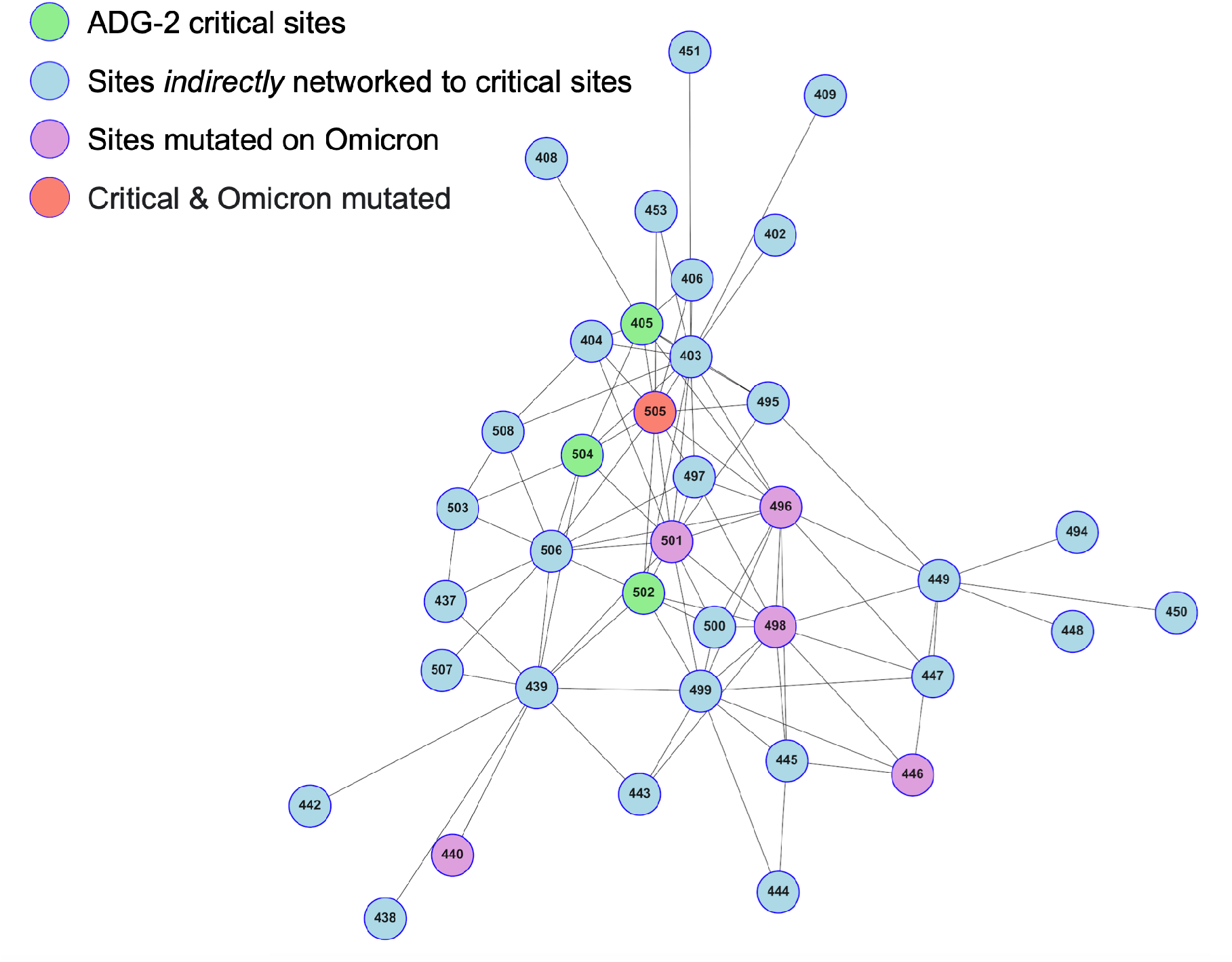
Indirect AAI network of the ADG-2 critical binding residues. RBD sites determined to be critical for ADG-2 binding [Rappazzo 2021] are shown in green or orange. RBD sites that indirectly interact with ADG-2 via the four critical sites are shown in blue and purple, where purple annotates sites mutated on the Omicron variant. Site 505 is shown as orange as it is both directly and indirectly networked to ADG-2. Edges between notes are weighted according to network interaction strengths, such that tightly clustered nodes with short edges between them are strongly networked.

